# Isolating soluble, near-native, misfolded states and their functional characterization: A step towards high-resolution protein misfolding structures

**DOI:** 10.64898/2026.11.24.769006

**Authors:** Ian Sitarik, Edward P. O’Brien

**Affiliations:** Institute for Computational and Data Sciences, Pennsylvania State University, University Park, Pennsylvania, United States; National Synthesis Center for the Emergence of Molecular and Cellular Sciences, Pennsylvania State University, University Park, Pennsylvania, United States; Department of Chemistry, Pennsylvania State University, University Park, Pennsylvania, United States; Bioinformatics and Genomics Graduate Program, The Huck Institutes of the Life Sciences, Pennsylvania State University, University Park, Pennsylvania, United States

## Abstract

Proteins can misfold into soluble, near-native states that act as long-lived kinetic traps with aggregation independent biological consequences. A key step toward obtaining high-resolution structures of these states is the ability to isolate/purify them. Here, we show this is possible for the *Escherichia coli* protein pyruvate kinase (*pykF*) following refolding from chemical denaturant. After controlling for aggregates the refolded (treatment) ensemble exhibits a decrease in average specific activity relative to the untreated (natively folded) sample. Size exclusion chromatography reveals two distinct populations in this treatment group, whereas the untreated group exhibits a single population. Native PAGE analysis finds that both populations from the treatment group have the same mass and electrophoretic mobility as the untreated protein, consistent with native-like global properties and oligomerization state. Functional analysis using Michaelis–Menten kinetics shows that one population retains activity comparable to the untreated sample, while the other has no detectable activity, indicating that approximately 31 ± 17.9% (95% Confidence Interval) of refolded molecules are soluble, near-native, but non-functional. ANS fluorescence measurements probing hydrophobic surface exposure further show that the non-functional population has altered surface properties relative to the active population. Differences in these properties persist for at least four hours following renaturation, indicating the system remains out-of-equilibrium. Together, these results demonstrate that size exclusion chromatography can be used to isolate soluble, non-aggregating, misfolded states that are structurally similar to the native state yet functionally inactive. This approach provides a basis for enriching such misfolded states for subsequent high-resolution structural characterization.

## Introduction

In a series of papers, we have provided evidence that soluble, long-lived, near-native misfolded states are a widespread phenomenon in vitro^1,2^ and also appears to occur in vivo^3,4^. This type of misfolding has gone unrecognized by biochemists primarily due to two prevailing assumptions that preclude such states from being common. First, proteins are generally expected to fold on biologically relevant timescales; therefore, if the native state is the global free-energy minimum under a given set of solution conditions, long-lived misfolded states are not expected to accumulate. Second, proteins that enter long-lived misfolded states are assumed to aggregate and partition into the insoluble fraction. Together, these assumptions leave little room for appreciable populations of soluble, long-lived, near-native misfolded proteins.

However, a recent meta-analysis by our laboratory of twenty experimental studies on the refolding of enzymes provided evidence that it is common for there to exist long-lived, soluble misfolded protein states that are non-functional^5^. The studies in the meta-analysis quantified the fraction of functional protein molecules versus time after a rapid switch from unfolding to refolding solution conditions. For the nine different proteins that were studied, the fraction of soluble, non-functional protein molecules ranged from 6 to 70% even in the presence of chaperones and most had estimated lifetimes of at least a day or longer. While many of the studies had controls to account for confounding factors, the conclusion that these misfolded states exist was instead inferred from kinetic fits and estimates of distinct protein subpopulations. The original studies never isolated and directly characterized the soluble, native-like misfolded molecules that supposedly exist.

Direct empirical confirmation of these subpopulations requires the isolation and purification of these highly native-like misfolded proteins, demonstrating they remain soluble over hours, and functionally characterizing them. This is a non-trivial task as many protein purification techniques require substantial changes in protein structure or surface properties^6–8^. Furthermore, such purification for the purposes of enrichment is a necessary first step in obtaining a high-resolution structure of these misfolded states^9,10^.

Here, we examine if it is possible to do this for the *E. coli* protein pyruvate kinase I (*pykF*) which was selected because Limited Proteolysis Mass Spectrometry (LiP-MS) experiments indicated this protein exhibits long-time-scale changes in protease sensitivity after being chemical unfolded and then refolded by dilution^2,11^. The protease digestion pattern changes reflect a structural change involving that misfolding in *pykF* that persisted for hours or longer. Thus, if soluble misfolded states exist, this is a promising candidate from which to attempt to purify them. Below, we show this can successfully be done with Size Exclusion Chromatography (Figure, 1).

**Figure 1.**
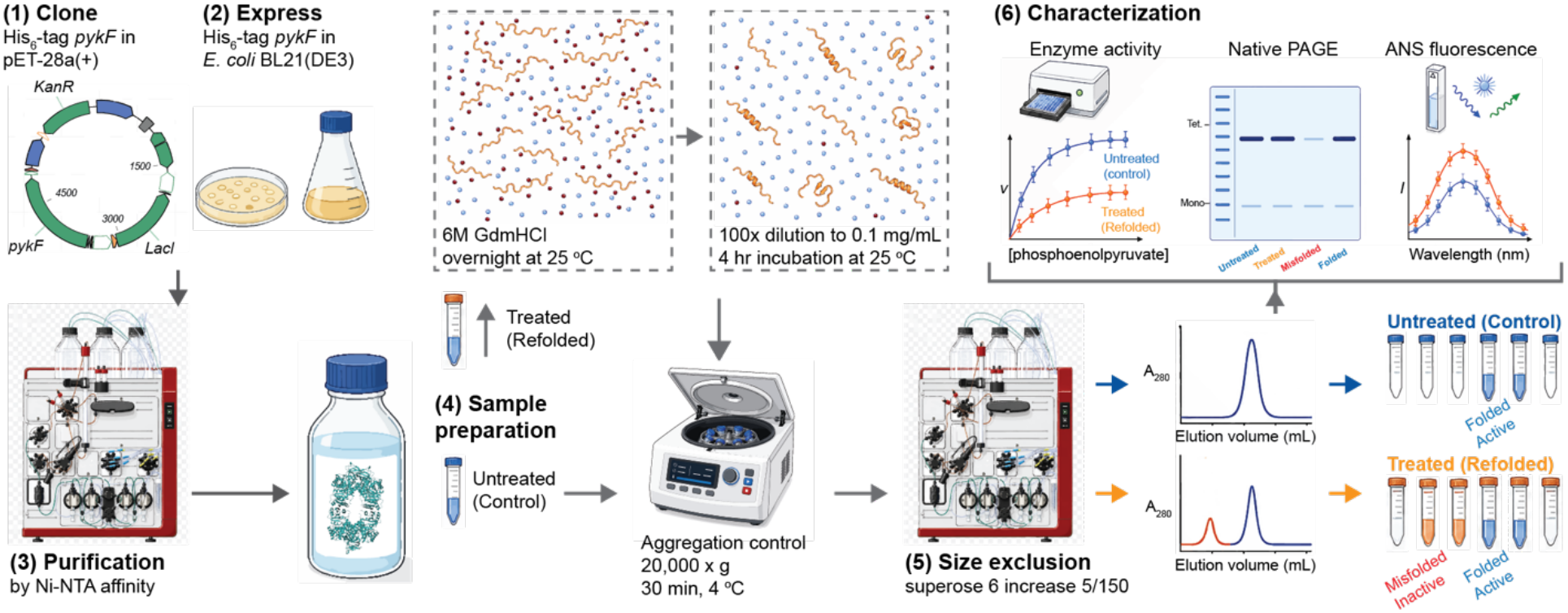
Experimental workflow for isolating and characterizing soluble, long-lived, near-native misfolded *pykF*. *pykF* was selected as a misfolding-prone candidate based on prior refolding and LiP-MS evidence, then cloned, expressed in *E. coli*, and purified by FPLC to obtain tetrameric protein in stable buffer. Purified *pykF* was split into untreated and refolding-treatment arms. The untreated control was carried directly into stable buffer, whereas the treated sample was unfolded in 6 M guanidinium HCl, diluted 100-fold into stable buffer to initiate refolding, and allowed to refold before clarification. Insoluble aggregates were removed by centrifugation, and the soluble supernatant was subjected to SEC-FPLC. Untreated *pykF* yielded a single predominant folded population, while refolded *pykF* separated into two soluble populations: an early-eluting near-native misfolded population and a later-eluting folded population. SEC-isolated fractions were then characterized by enzyme activity assays, native PAGE, and ANS fluorescence to assess catalytic function, oligomeric state, and exposed hydrophobic surface area.

## Results

### A collection of soluble, pyruvate kinase molecules change their average specific activity upon refolding that persists for hours

If *pykF* has subpopulations of molecules that misfold upon sample wide refolding, as suggested by LiP-MS, ensemble level measures of function should also decrease. To test this, we treated a sample of *pykF* with 6 M guanidinium hydrochloride for 12 hours to induce unfolding and established refolding conditions through dilution to 0.06 M and measured the resulting activity of the sample. We carried out initial velocity experiments four hours after refolding conditions were established, measured the Michaelis-Menten parameters and Hill coefficient^12^ (a measure of cooperativity), and find that between the untreated and guanidinium-treated samples, the *k*_*cat*_ is 30% lower in the treated sample (Table 1 untreated *k*_*cat*_ = 101.5 ± 19.58 s^-1^ ; treated *k*_*cat*_ = 70.07 ± 12.09 s^-1^; difference *p* = 1 x 10^-5^, 3 biological replicates, permutation test). However, the Michaelis-Menten constant *k*_*M*_ and the Hill coefficient *n* are unchanged (Figure, 2a; Table 1). Thus, we conclude that upon refolding there is a significant alteration in the activity of *pykF* despite the same concentration of soluble protein being present. This result is consistent with one or more subpopulations of *pykF* misfolding and subsequently altering the overall activity of the collection of *pykF* molecules.

**Table 1.** Michaelis Menten parameters of untreated and refolded (treated) pyruvate kinase.

|  | *k*<sub>*cat*</sub>, 1/s | *K*<sub>M</sub>, mM | *n* |
|---|---|---|---|
| Untreated PK | 101.5 ± 19.58 | 7.12 ± 0.48 | 4.67 ± 1.15 |
| Treated PK (2 mL + 1 mL elution volumes) | 70.07 ± 12.09 | 7.57 ± 0.64 | 4.42 ± 1.20 |
| Untreated folded peak (2 mL elution volume) | 99.80 ± 11.81 | - | - |
| Treated folded peak (2 mL elution volume) | 105.12 ± 12.98 | - | - |
| Treated misfolded peak (1 mL elution volume) | 0.171 ± 1.88 | - | - |

### Soluble *pykF* molecules split into two distinct populations

If there is indeed a mixture of soluble misfolded and folded *pykF*, the misfolded states may either have changes in their surface properties (i.e., increased exposure of hydrophobic surface area) or changes in their oligomeric structure making them susceptible to separation by liquid chromatography. To experimentally test this hypothesis, we applied size-exclusion fast protein liquid chromatography (SEC-FPLC) on both the untreated and treated samples. We observed one population in the untreated sample at 2 mL of elution volume (Figure, 2b), which we presume is made up of mostly folded *pykF* molecules. In the treated sample we still observe a folded population at 2 mL elution volume, however, a new population also appears at 1 mL elution volume (Figure, 2b). We conclude that four hours after establishing folding conditions there is a population of *pykF* molecules that adopt the native conformation and an additional population of soluble *pykF* that is structurally distinct.

### The two populations appear folded in Native PAGE

Native *pyfK* is predominantly a tetramer^13^ and therefore we hypothesized that this new subpopulation could be due to a change in the distribution of oligomeric states upon refolding rather than forming a soluble non-functional tetramer. To determine this, we ran native (non-denaturing) PAGE gels, which preserve the oligomerization state, of (*i*) the untreated sample, (*ii*) the treated sample, (*iii*) the folded population SEC peak (i.e., 2 mL elution volume) of the treated sample, and (*iv*) of the 1 mL elution volume population of the treated sample. In all four gels we observe one predominant band, corresponding to the molecular weight of a *pyfK* tetramer (Figure, 2c), and a weaker band corresponding to *pyfK* monomer. Thus, we conclude that in the treated sample, and the two subpopulations that compose it, the distribution of oligomers is consistent with the untreated sample and hence consistent with forming soluble, native-like tetramers.

### One population is as functional as the untreated sample, the other has no activity

Next, we characterized the functionality of each population. From the SEC-FPLC, we collected fractions from each population in the treated sample as well as from the single peak in the untreated sample and measured the *k*_cat_ under saturating conditions. For the folded population peak in both the treated and untreated samples we observe they have statistically indistinguishable values (Table 1 treated folded peak *k*_cat_ = 105.12 ± 12.98 s^-1^ ; Table 1 untreated folded peak 99.80 ± 11.81 s^-1^, *p* = 0.99, 8 technical replicates, permutation test). Thus, this SEC-FPLC peak in the treated sample corresponds to folded, functional *pykF*. In contrast, the treated sample’s additional population (elution volume of 1 mL) has essentially no activity with *k*_cat_ = 0.171 ± 1.88 s^-1^ (see Figure 2b and Table 1 treated misfolded peak). Thus, we conclude that upon refolding, *pykF* partitions into two populations: a population of folded native and active tetramers and a population of misfolded tetramer molecules that are kinetically trapped (i.e., have not achieved thermodynamic equilibrium) in a soluble form for at least four hours.

**Figure 2.**
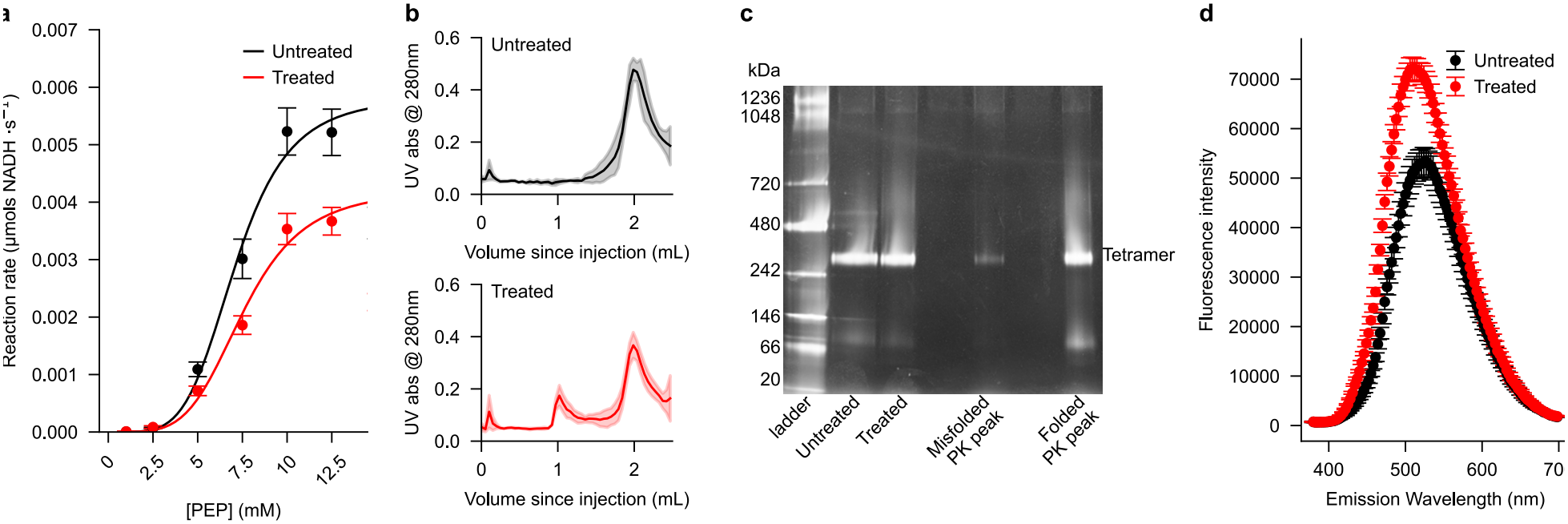
Refolding of *pykF* produces soluble, misfolded tetrameric states with reduced activity. **(a)** Initial velocity of *pykF* as a function of phosphoenolpyruvate (PEP) concentration for untreated and refolded samples across three biological replicates. Refolded samples were unfolded in 6 M guanidinium chloride and returned to folding conditions by dilution. Data were fit to the Hill equation^12,45^ to estimate apparent *k*_*cat*_, *K*_*M*_, and Hill coefficient *n*. Refolded *pykF* shows reduced catalytic activity, while *K*_*M*_and cooperativity remain largely unchanged. **(b)** Average SEC-FPLC elution profiles of untreated and refolded *pykF* after centrifugation at 20,000 × *g*, monitored at 280 nm. Untreated samples show a folded population at approximately 2 mL, whereas refolded samples contain this population and an additional earlier-eluting soluble population at approximately 1 mL. Shading represents 95% confidence intervals across biological replicates. **(c)** Blue native PAGE of untreated and refolded *pykF* and SEC-FPLC peak fractions from the refolded sample. Lanes 2 and 3 show total soluble samples; lanes 5 and 7 show the earlier-eluting perturbed population and later-eluting folded population from panel b. Both migrate consistently with tetrameric *pykF*. **(d)** ANS fluorescence emission spectra of untreated and refolded *pykF* after excitation at 365 nm. Refolded *pykF* shows increased ANS fluorescence, consistent with increased hydrophobic surface exposure. Shading represents 95% confidence intervals from technical replicates.

**Figure 3.**
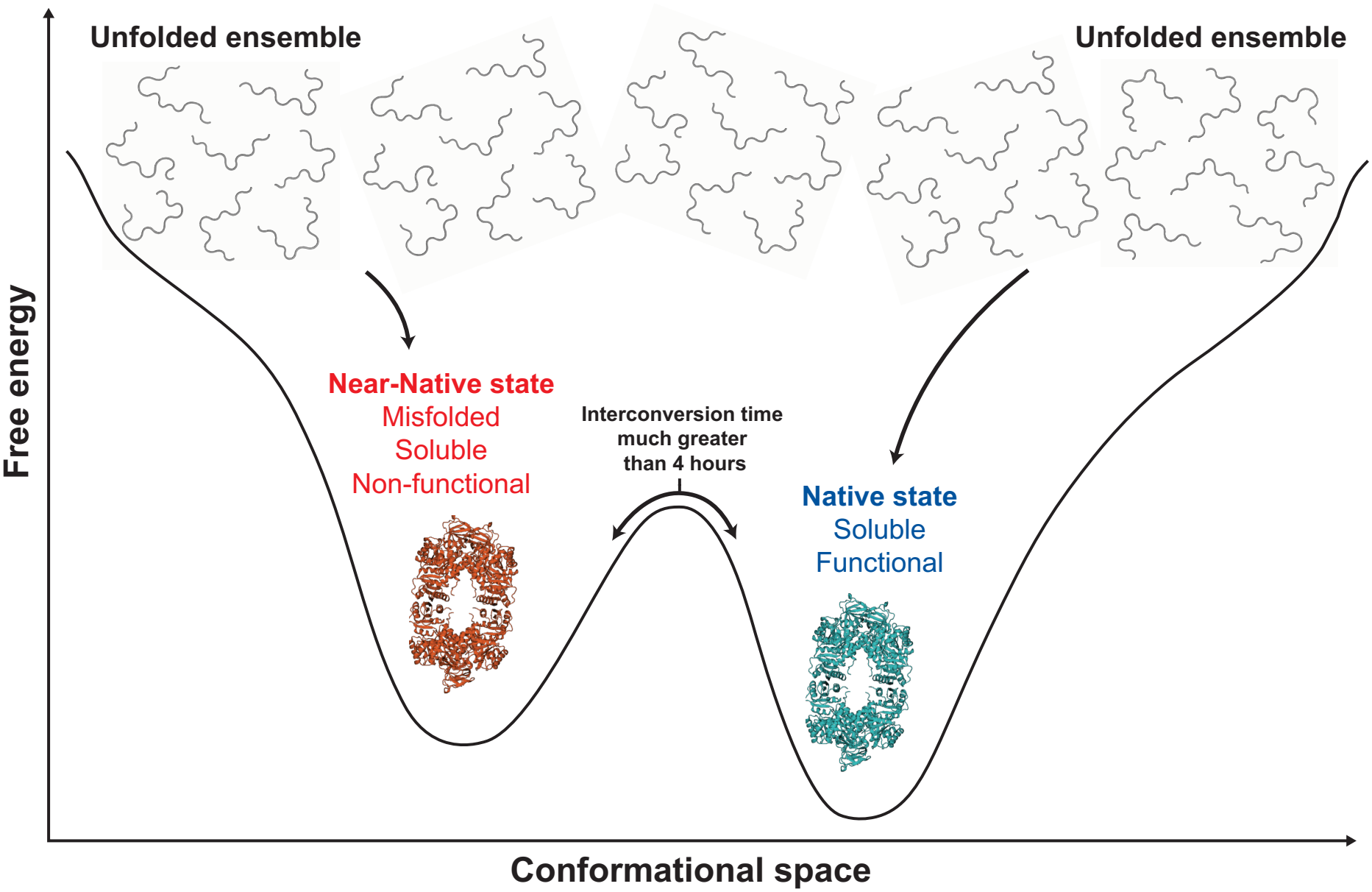
Energy landscape for kinetic trapping of soluble, long-lived, near-native misfolded *pykF*. Our data supports a folding-energy landscape in which refolded *pykF* can partition into distinct soluble tetrameric states, rather than proceeding rapidly and uniformly to the native ensemble. Upon dilution from denaturing conditions, a subset of molecules reaches the native state, which is functional and assumed here to represent the global free-energy minimum (right-hand side of landscape). A second subset of molecules becomes trapped in a local free-energy minimum corresponding to a soluble, near-native misfolded state (left-hand side of landscape). This state retains native-like global structure and oligomeric organization but is non-functional, consistent with its separation by SEC-FPLC, tetramer-like mobility by native PAGE, loss of catalytic activity, and altered surface exposure by ANS fluorescence. The model illustrates how soluble misfolded proteins can persist as kinetically trapped molecular states despite the native state presumably being thermodynamically favored. Note well, there must be a large free energy barrier between the states as the interconversion time between these two states is much greater than 4 hours in our experiments. This free energy surface is qualitatively consistent with the Kinetic Partitioning Mechanism^19^.

### The treated sample has increased hydrophobic surface area

Since the 1 mL elution volume subpopulation is still forming soluble *pykF* tetramer (Figure, 2c) that is non-functional (Table 1) and has decreased interactions with the hydrophilic SEC-FPLC column (as it runs faster through the column), we hypothesized that this misfolded population must have an increase in exposed hydrophobic surface area. To test this hypothesis, we applied an 8-Anilino-1-naphthalenesulfonic acid (ANS) assay, which fluoresces when bound to hydrophobic residues^14,15^, to the treated and untreated samples with identical concentrations of *pykF* molecules. We observe a large and significant increase in the emission intensity in the treated sample relative to the untreated sample, indicating increased exposure of hydrophobic residues in the misfolded tetramer, but not apparently enough to cause insoluble aggregation over the four-hour time course of this study (Figure, 2d). Therefore, we conclude that upon refolding, a population of *pykF* molecules structurally misfold into near native-like structures that persist for hours or longer.

## Discussion

We have provided direct experimental confirmation of the existence of soluble, long-lived, near-native, misfolded protein states; specifically, we demonstrated that they can be physically isolated and functionally characterized. We found that the refolding of *E. coli pykF* produces two distinct soluble populations: one that is fully functional and one that is inactive. These populations are separable by size exclusion chromatography (SEC), despite exhibiting indistinguishable oligomeric states by native PAGE and comparable global electrophoretic mobility. The misfolded population remains soluble following ultracentrifugation, persists for at least four hours after refolding, and displays altered surface hydrophobicity as measured by ANS fluorescence. Together, these observations establish a two-state partitioning of the refolded ensemble into native and misfolded basins that persist on experimentally relevant timescales.

Several aspects of these results are notable. First, we ruled out trivial explanations for the observed loss of activity. The absence of changes in oligomeric state (native PAGE) eliminates dissociation or aggregation-driven changes^16,17^. Focusing on the supernatant after ultracentrifugation, which separates insoluble and soluble molecules^18^, demonstrates that the inactive population we characterized is not insoluble or aggregated. The persistence of the misfolded population at the four-hour time point indicates that these are long-lived kinetic traps rather than transient folding intermediates, and is consistent with an energy landscape described by the Kinetic Partitioning Mechanism^19^ (Figure, 3). Finally, the clean functional separation into fully active and fully inactive populations supports a discrete, two-state description at the level of functional output, even if underlying structural heterogeneity exists.

The ability to directly isolate and characterize these states has important implications for cellular proteostasis. Prior evidence for soluble misfolded states has been indirect^2,3,5^ - based on ensemble measurements and inference after excluding alternative explanations. Here, by physically separating the inactive population, we provide direct confirmation that a substantial fraction (∼30% in this system) of the protein molecules occupy a soluble, non-functional state. In cellular contexts, such states may be particularly consequential: if sufficiently native-like, they have been suggested to evade the chaperone and degradation quality control pathways that target gross misfolding^1,2,4,5^. This suggests that a significant population of proteins in vivo likely exist in “zombie” misfolded states, that have no function but are native-like enough to bypass chaperone recognition and avoid rapid aggregation and degradation. We therefore conclude that these states constitute a previously unrecognized component of the proteostasis network^4^ that is distinct from the canonical categories of folded, unfolded, aggregated or degraded proteins.

To our knowledge, there are relatively few precedents for physically separating a non-functional, soluble subpopulation of a protein that is otherwise native-like in size and oligomerization state. Classical approaches to alternative conformational states such as ion exchange chromatography^6^, ligand-based selection^7^, or temperature/pH-induced^8^ conformational trapping, typically enrich for states that differ substantially in charge, stability, or folding cooperativity. More recently, kinetic partitioning studies from our lab and others have indicated the existence of long-lived, non-native states using functional or kinetic readouts^3,20^, but these approaches do not physically isolate the inactive population for direct biochemical characterization. In contrast, our method leverages SEC to resolve populations that are similar in mass and oligomeric state but differ subtly in surface interactions with the stationary phase, enabling both purification and downstream functional assays. This represents a conceptual and technical advance: rather than inferring misfolded states from ensemble behavior, we isolate them as discrete biochemical entities.

The fine structural details of these misfolded states is an important question that we did not address in this study. Two plausible models are domain swapping^21–23^ and changes in non-covalent lasso entanglements (NCLEs). Domain swapping is a mechanism in which structural elements are exchanged between subunits that are similar in structure and sequence, producing oligomers that retain native-like features but are topologically altered^23–25^. Such rearrangements could preserve tetrameric assembly and gross structure while disrupting catalytic function. Alternatively, NCLEs are structural motifs in which a segment of the polypeptide threads through a loop stabilized by non-covalent interactions, providing a topological mechanism for trapping proteins in near-native conformations that are kinetically difficult to resolve^2,20^. Unlike domain swapping, NCLEs do not require subunit exchange and can arise within a single polypeptide chain and their association with highly non-refoldable proteins has been established^2^. Additionally, non-native knot formation has also been proposed as a misfolding mechanism for *E. coli* phosphoglycerate kinase^26^. The current study does not establish which folding mechanism is occurring in *pykF*, but it enables future studies to resolve this through greater structural resolution of the isolated misfolded ensemble. Importantly, disulfide bond formation is unlikely to explain the observed behavior, as the experimental conditions include reducing agents (β-mercaptoethanol), which makes disulfide bond formation unfavorable^27^, and the native fold of *pykF* lacks disulfide bonds.

While domain swapping has been extensively characterized structurally, NCLE-mediated misfolding has yet to be captured at high resolution. The present work therefore provides a critical first step: by isolating and enriching the misfolded population, we create the opportunity for direct structural interrogation. Notably, the separation achieved by SEC reflects not only differences in hydrodynamic radius but also secondary, often called ‘non-ideal’, contributions from protein– matrix interactions^28^. The earlier elution of the misfolded population suggests reduced interaction with the column matrix, Sepharose (a polar media), consistent with increased exposure of hydrophobic surface area. The increase in hydrophobic surface area was independently supported by ANS fluorescence changes. Thus, SEC in this context acts as a hybrid separation method, sensitive to not only size, a primary factor, but also the secondary factor of surface chemistry.

To obtain high-resolution structures using Cryo-EM higher concentrations of misfolded protein are needed than what we obtain in the SEC eluent in this study. There are a number of biochemical techniques that can increase the concentration while preserving conformational states, such as osmotic dialysis^29^. Together, these approaches offer a path toward resolving the structural basis of these misfolded states and determining whether mechanisms such as those involving NCLEs are operative.

In summary, we demonstrated that soluble, near-native, misfolded protein states can be directly isolated and functionally characterized. These findings, consistent with a number of recent studies^1–3,5,20,30–32^, challenge the conventional view that long-lived misfolding is tightly coupled to aggregation and instead point to a substantial and potentially biologically relevant population of kinetically trapped, soluble states. By enabling their purification, this work lays the foundation for future high-resolution structural studies and opens a new avenue for understanding protein misfolding within the broader landscape of proteostasis.

## Methods

### Candidate selection

Pyruvate kinase activity has been extensively studied for 30+ years^33–44^ and in some organisms is a curcial rate limiting step in glycolysis. It is often used as a “work-horse enzyme” in a coupled enzyme assay with lactate dehydrogenase to measure the production of ADP by a target enzyme. It was also previously identified to be a protein that has trouble refolding on long time scales by limited proteolysis mass spectrometry and it has a unique misfolding prone structural motif called a native non-covalent lasso entanglement which has a thread that takes part in the active site. Therefore it makes a prime candidate for investigation and separation.

### Cloning and overexpression of pyruvate kinase I in E. coli

The canonical DNA sequence for pyruvate kinase I was pulled from Uniprot (P0AD61) and the fragment was synthesized by Genescript for cloning into a pET-26b(+) vector (New England Biolabs) containing a C-terminal hexahistidine (His6) tag using Ndel and SacI restriction sites. The resulting construct, pET-26b(+)-PK was verified by DNA sequencing and then used to transform E. coli BL21 (DE3) competent cells (ThermoFisher). Colonies plates were grown overnight on media consisting of lysogeny broth (LB) agar mixture w/ 50mg/L Kanamycin. Single colonies were selected to inoculate 50mL of prewarmed (37°C) LB starter culture containing 50mg/L Kanamycin followed by incubation at 37 °C and shaking 180 rpm for 12 hours. 5mL of the starter culture was then used to inoculate 1L of LB medium containing 50mg/L kanamycin which was incubated at 37 °C and 180 rpm until an optical density at 600nm (OD600) of ∼0.6 was reached. Protein expression was then induced by the addition of 0.25mM isopropylthio-β-D-galactoside (IPTG), and incubation was continued at 25 °C at 180 rpm for an additional 12-15 h. The cells were harvested by centrifugation (4 °C, 15000 × g, 15 min) and flash frozen in 2g chunks in N2(l) followed by storage at -80 °C. Samples were labeled with E#P# where E# indicates the biological replicate and P# indicates which pellet of that replicate was used. Not all pellets were used.

### Purification of pyruvate kinase I from E. coli

The cells were resuspended in lysis buffer (20 mM Tris-HCl, 200 mM NaCl, 50 mM MgSO4, 100 mM KCl, 5 mM imidazole, 2 mM β-mercaptoethanol pH 7.5) at 4 °C and on a rotor at 60 rmp for 40 min. The mixture was then lysed by sonication with an ultrasonic cell disruptor (Branson 450) in an ice water bath with x10 30-second pulses with a power level of 6 and a duty-cycle of 60% and a 1 min rest period between pulses. The lysate was then clarified by centrifugation (4 °C, 45000 × g, 45 min). The supernatant was then vacuum filtered over 0.22µm membrane filters (EZ-Pak Millipore) at 4 °C. All following purification steps were done in a cold room at 4 °C. The filtrate was then loaded onto a XK-16/20 Cytiva column with a peristaltic pump filled with 15 mL Ni-Nitrilotriacetate resin (Ni-NTA, HisPur ThermoScientific) that was pre-equilibrated in lysis buffer. The resin was then washed at 1 mL/min with lysis buffer till the UV280nm trace reaches a baseline followed by a wash with wash A (20 mM Tris-HCl, 200 mM NaCl, 50 mM MgSO4, 100 mM KCl, 20 mM imidazole, 2 mM β-mercaptoethanol pH 7.5) at 1 mL/min till the UV280nm trace reaches a baseline followed by a wash with wash B (20 mM Tris-HCl, 200 mM NaCl, 50 mM MgSO4, 100 mM KCl, 40 mM imidazole, 2 mM β-mercaptoethanol pH 7.5) at 1 mL/min till the UV280nm trace reaches a baseline. Finally, the bound protein was eluted from the column with (20 mM Tris-HCl, 200 mM NaCl, 50 mM MgSO4, 100 mM KCl, 300 mM imidazole, 2 mM β-mercaptoethanol pH 7.5). The collected fractions were then concentrated with 30kDa MWCO viva-spin filters down to 1mL (1000 × g) and then buffer exchanged into stable buffer (20 mM Tris-HCl, 200 mM NaCl, 50 mM MgSO4, 100 mM KCl, 2 mM β-mercaptoethanol pH 7.5) with Cytiva HiTrap desalting column (5 mL) pre-equilibrated in stable buffer. The final sample was then concentrated to ∼35 mg/mL judged by UV absorption at 280nm and stored for no more than 1 week at 4 °C. Column resin was cleaned with the following series of buffers: 5 column volumes (CV) of purified water, 5CV of 20% EtOH, 5CV of 20 mM 2-(N-morpholino)ethanesulfonic acid (MES) buffer, 5CV of purified water, 200 mM NiSO4, 5CV of purified water, and finally stored in 20% EtOH till next use.

### Preparing refolded samples of pyruvate kinase I from E. coli

Samples were selected at random from the available pellets for a given biological replicate. Treated sample described in this work were prepared by unfolding PK by dissolving 28.6 mg of Guanidine Hydrochloride (Millipore sigma, Ultrol® grade) in 35 uL of stable buffer (20 mM Tris-HCl, 200 mM NaCl, 50 mM MgSO4, 100 mM KCl, 2 mM β-mercaptoethanol pH 7.5) and then adding 15 uL of purified PK sample at 33.3 mg/mL yielding a 6M denaturing environment^11,36^. The sample was unfolded overnight at 25 °C and then diluted 100x by delivery into 4950 uL of stable buffer yielding a 0.1 mg/mL PK refolding environment. This is the maximal refolding concentration where we do not observe visible aggregation. Mixing was done with a pipette to minimize bubbles formation and allowed to refold for 4 hours on the at 25 °C. To control for aggregation we centrifuge the sample at 20,000 × g for 30 min at 4 °C before any further experiments are done. Untreated samples were prepared in the same way except the PK is not added till after the 100x dilution step.

### Size exclusion chromatography

A Cytiva Superose™6 increase 5/150 GL column was pre-equilibrated in stable buffer (20 mM Tris-HCl, 200 mM NaCl, 50 mM MgSO4, 100 mM KCl, 2 mM β-mercaptoethanol pH 7.5) with 20 CV at 0.3 mL/min and 4 °C. Samples were loaded into a 100uL sample loop that was pre-washed with stable buffer and then injected onto the column and run through at 0.3 mL/min and 4 °C. In-between samples the column was washed with 10 CV of purified water and re-equilibrated in stable buffer. After 5 samples were run the column was cleaned with 10 CV 20% EtOH.

For peak selection to determine population activity in saturating substrate conditions and oligomeric states we first concentrated the refolded or control samples in a 30kDa MWCO viva-spin filter (500 × g) that were pretreated with stable buffer (20 mM Tris-HCl, 200 mM NaCl, 50 mM MgSO4, 100 mM KCl, 2 mM β-mercaptoethanol pH 7.5, incubate 2 hours at 25 °C) and then centrifugation at 3000 × g for 30 min at 4 °C. Peak fractions were taken in 150uL samples and those closest to the peak center were used in all subsequent experiments to avoid dilution effects and cross population contamination.

### Michaelis–Menten kinetics

Enzyme kinetics was measured using previously described procedures with a Spectramax Plus 384 UV-vis plate reader and some minor modifications. Reactions contained the following in a final volume of 300 uL: 45 mM Imidazole, 110 mM KCl, 55 mM MgSO4, 2 mM ADP, ∼0.0125 mg/mL PK, 0.12 mg/mL lactate dehydrogenase, 0.9 mM NADH, and varying concentrations of phophoenol pyruvate (PEP, 1 – 20 mM)^35,37,39,45^. Reaction was initiated by the addition of 25 uL of PEP and conversion of NADH was monitored over 5 min at 37 °C by the loss in absorbance at 340 nm (ε340=6440 M-1 cm-1). Technical replicates (n = 8) were done for each concentration of PEP for each biological replicate (n = 3). Initial velocities were extracted by linear fitting the first 30s of each kinetic trace for [PEP] = 20, 17.5, 15, 5, 2.5, 1 mM and the first 15 s for [PEP] = 12.5, 10, 7.5 mM. The resulting data was fit using the Hill equation^12,45^ to estimate the rate constant (K_cat_), the apparent Michaelis-Menten constant (K_M_), and the Hill coefficient (n)^12^. Confidence intervals were calculated assuming a normal distribution.

### SDS and Native PAGE

For denaturing sodium dodecyl sulfate–polyacrylamide gel electrophoresis (SDS PAGE) 10 uL of each sample was added to 10 uL of sample buffer (Novex™ Tris-Glycine SDS sample buffer 2x). If DTT was required 0.5 uL of 1M DTT was added and then samples were heated for 10 min at 80 °C. Sample volumes of 20 uL were then loaded into each well of the gel (Novex™ Wedgewell 12% Tris-Glycine) and the chambers filled with running buffer (Sigma-Aldrich Tris-glycine SDS buffer 1x). SDS-PAGE was run at 25 °C for 40 min at 225 V.

For Native PAGE 10 uL of sample was mixed with 10 uL of loading buffer (Novex™ Tris-Glycine Native Sample Buffer (2X)) and let sit at 4 °C for 10 min. Sample volumes of 10 uL were then loaded into Invitrogen NativePAGE™ 3-12% Bis-Tris gel wells with Invitrogen NativeMark™ unstained protein ladder as a standard reference. 200 mL of NativePAGE™ light blue cathode buffer (1x) was added to the inner chamber and 600 mL of NativePAGE™ anode buffer (1x) was added to the outer chamber. PAGE was then run at a constant 150 V for 60 min and then 250 V for 30 min at 4 °C as per the manufacturer protocol.

Silver staining both SDS and Native PAGE gels was done in the following steps^46^. The gel was first incubated in 50% MetOH for 15 min (on a rocker table at 25 °C), then 5% MetOH for 15 min, and then 32 uM DTT for 15 min. The gel was then briefly washed twice in purified water and incubated in 0.1% AgNO3 for 15 min. The gel was then briefly washed with purified water for 1 min. The developing solution (3% Na2CO3) was made fresh and 2 min before use 50 uL formaldehyde was added to 100 mL of the solution and mixed. The gel was then incubated in the activated developing solution until the desired band intensity was observed and then imaged.

### 8-Anilino-1-naphthalenesulfonic acid assay for changes in hydrophobic exposure

Changes in the extent of hydrophobic exposure were probed by changes in the fluorescence intensity of 8-Anilino-1-naphthalenesulfonic acid (ANS)^14,15,47^. Technical replicates (n = 8) each contained 250 uL of: ∼ 2 uM PK samples (refolded or untreated), 0.5 mM ANS, 20 mM Tris-HCl, 200 mM NaCl, 50 mM MgSO4, 100 mM KCl, 2 mM β-mercaptoethanol at pH 7.5. Samples were allowed to equilibrate for 1 hour at 25 °C before excitation at 365 nm and measuring emission from 380 – 700 nm. Confidence intervals at each emission wavelength were calculated assuming a normal distribution.

### Sample concentrations estimated by Bradford

Sample concentrations for all experiments after purification were estimated by Bradford calibration as described in the Pirce™ Bradford Plus Protein Assay Kit (Thermo Fisher) with the following modifications. The standard used was our purified pyruvate kinase instead of the typical BSA. We used 20 uL of sample mixed with 20 uL of Coomassie Plue reagent incubated for 10 min at 25 °C before 2 uL was examined for absorption on the Nanodrop (Thermo Scientific). Calibration curve.

## Acknowledgements

E.O. gratefully acknowledges support from the National Science Foundation (MCB-2031584) as well as from the National Institutes of Health (R35-GM124818). Portions of numerical computations and data analysis in this work have been carried out on high-performance computing collectively known as ROAR, which is operated by the Institute for Computational and Data Sciences at The Pennsylvania State University. This work was supported by the National Science Foundation (NSF) through the NSF National Synthesis Center for Emergence in the Molecular and Cellular Sciences (NCEMS) under Grant NSF MCB-2335029.

## Credit authorship contribution statement

**I.S**.: Methodology development, Experimental data collection, Formal analysis, Visualization, Validation, Data curation, Writing – original draft, Writing – review and editing.

**E. O**.: Conceptualization, Formal analysis, Funding acquisition, Investigation, Methodology development, Project administration, Resources, Supervision, Validation, Writing – original draft, Writing – review and editing.

