## Supplemental Information for "Isolating soluble, near-native, misfolded states and their functional characterization: A step towards high-resolution protein misfolding structures"

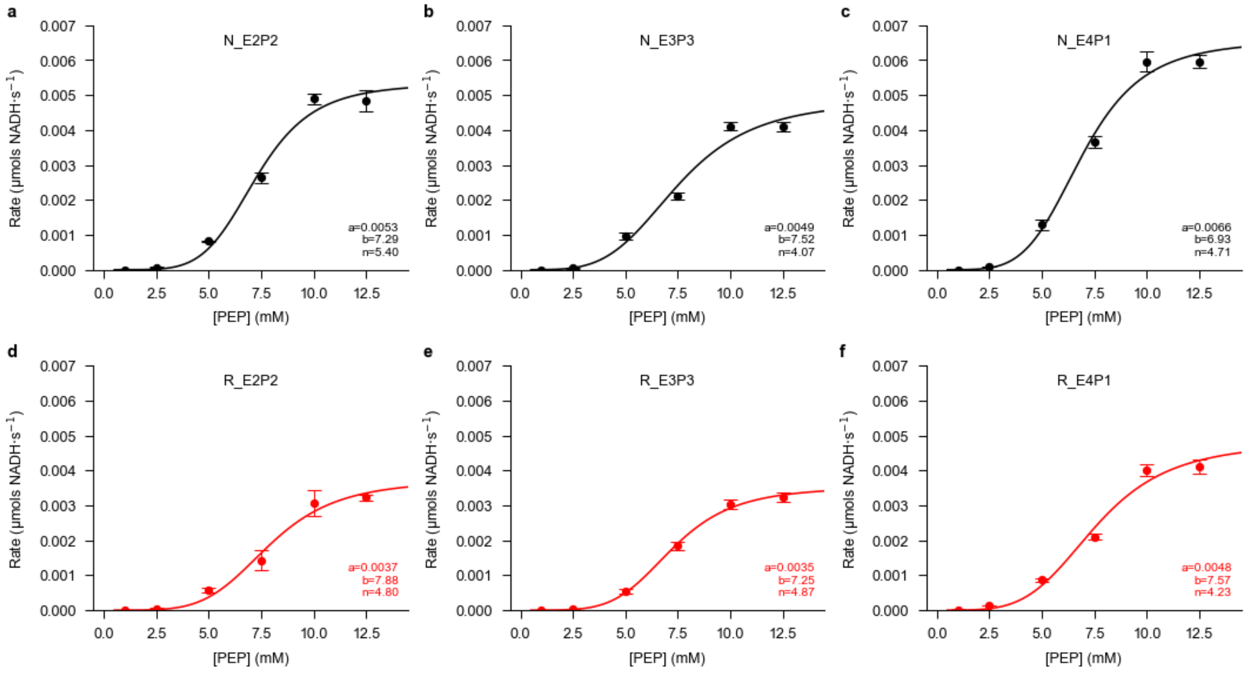


**Figure S1. *pykF* reaction kinetics for each biological replicate in treated and untreated samples.** (a–c) Initial velocities of untreated *pykF* plotted as a function of phosphoenolpyruvate (PEP) concentration for each of three biological replicates. (d–f) Initial velocities of *pykF* measured 4 h after unfolding in 6 M guanidinium chloride followed by refolding by 100x dilution. Reactions were monitored by NADH consumption at 340 nm in a coupled lactate dehydrogenase assay at 37 °C. For each PEP concentration, technical replicate traces were used to estimate initial velocities, which were then fit to the Hill equation to determine apparent kinetic parameters, including $k_{cat}$, $K_M$, and the Hill coefficient $n$. Points represent technical replicate measurements and curves show the best-fit Hill model for each biological replicate. The treated samples show reduced apparent catalytic activity relative to untreated *pykF*, while maintaining similar apparent $K_M$ and cooperativity. Samples were labeled with “N_” for untreated and “R_” for the refolded while E# dictates the biological replicate and P# indicates which pellet of that replicate was used.

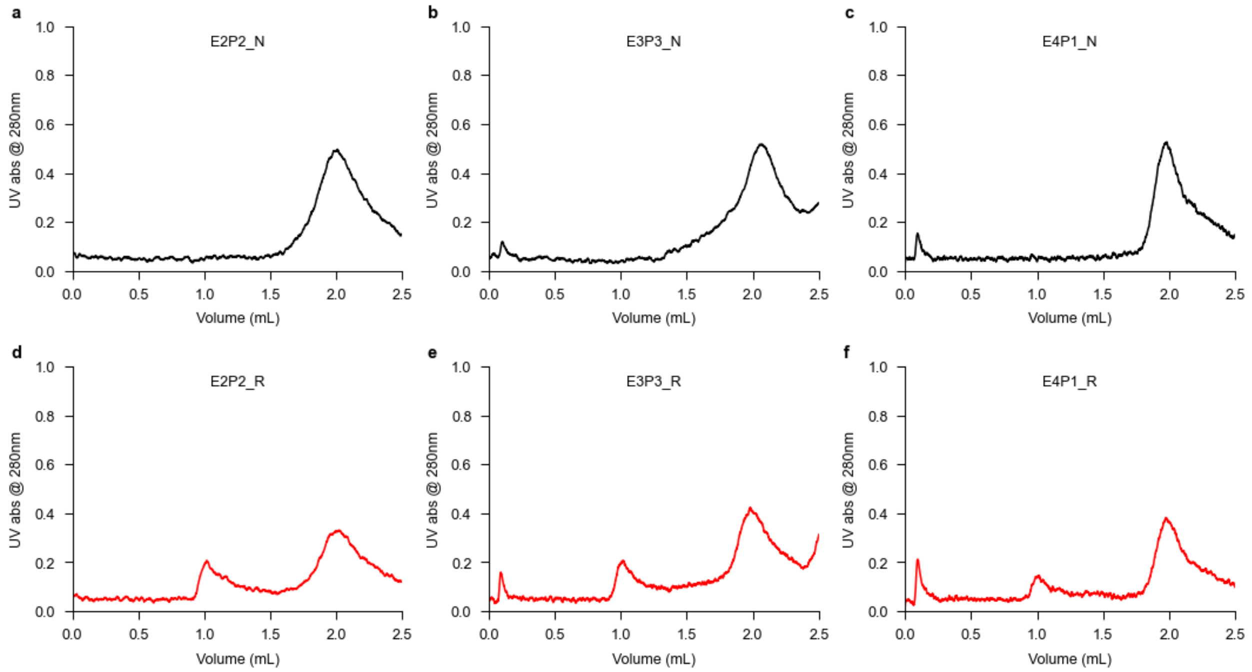


**Figure S2. *pykF* size exclusion chromatography (SEC) elution profiles for treated and untreated samples across biological replicates.** (a–c) Size-exclusion FPLC chromatograms of untreated pykF for three biological replicates following clarification by centrifugation at 20,000 × g. Protein elution was monitored by absorbance at 280 nm. Untreated samples show a dominant peak at approximately 2 mL elution volume, corresponding to the soluble folded *pykF* population. The small feature near 0.15 mL is a column-associated artifact also observed in blank runs. (d–f) SEC-FPLC chromatograms of *pykF* samples measured 4 h after unfolding in 6 M guanidinium hydrochloride followed by refolding by dilution. In addition to the folded population at approximately 2 mL, treated samples contain a distinct earlier-eluting soluble population at approximately 1 mL. This earlier-eluting peak represents a structurally perturbed *pykF* population that is separable from the folded population under native SEC-FPLC conditions. Samples were run on a Superose 6 Increase 5/150 GL column at 4 °C in stable buffer, and peak-centered fractions were collected for subsequent biochemical analysis. Samples were labeled with “N_” for untreated and “R_” for the refolded while E# dictates the biological replicate and P# indicates which pellet of that replicate was used.

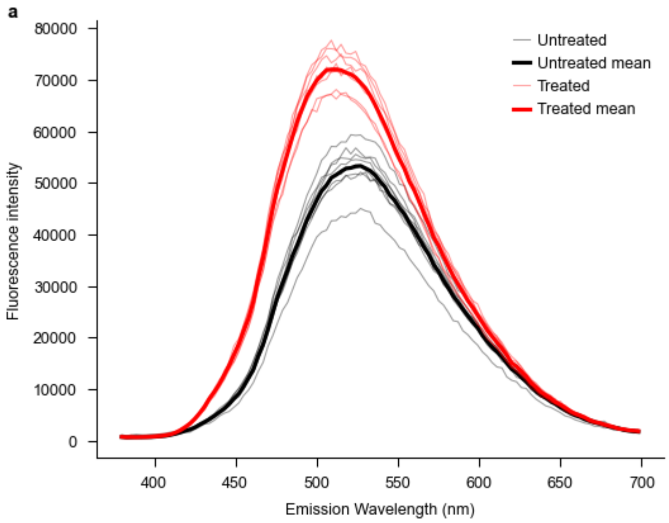


**Figure S3. *pykF* ANS fluorescence for treated and untreated samples across biological and technical replicates.** ANS fluorescence emission spectra for untreated *pykF* and *pykF* measured 4 h after unfolding in 6 M guanidinium chloride followed by refolding by dilution. Samples contained matched concentrations of soluble *pykF* and were incubated with 0.5 mM 8-anilino-1-naphthalenesulfonic acid (ANS) for 1 h at 25 °C before excitation at 365 nm. Emission was monitored from 380–700 nm. Refolded *pykF* samples show increased ANS fluorescence relative to untreated *pykF*, consistent with greater exposure of hydrophobic surface area in the structurally perturbed soluble population. Curves represent biological replicates, with technical replicates used to estimate uncertainty at each emission wavelength.
